# Use of solvent extraction–indigenous microbial degradation technology to repair soils contaminated by high concentrations of mechanical oil

**DOI:** 10.1101/664490

**Authors:** Guo Chen, Zhongyi Yin, Anping Liu, Xuxu Zheng

## Abstract

Remediation of soils contaminated by mechanical oil has become a difficult problem worldwide. In this study, soil contaminated by mechanical oil was repaired by domestication and inoculation of microorganisms collected from the contaminated site, and soil quality and plant growth indexes were evaluated to verify the efficacy of our solvent extraction–indigenous microbial degradation technology. Solvent extraction of the contaminated soil removed 97.03% mechanical oil, increased soil water-holding capacity by 68.20%, and improved root activity and soluble sugar content of alfalfa plants. However, solvent extraction depleted 82.98% of the soil organic matter. Screening and domestication of strain TB-6 from the contaminated site revealed that it is an *Enterobacter* with obvious degradation effects on petroleum hydrocarbons (C15–C28). After the solvent-extracted soil was inoculated with TB-6 for 30 days, the soil structure became loose; degradation rate of residual mechanical oil was 19.45%; and soil organic matter content, water-holding capacity, alfalfa root activity, and soluble sugar content increased by 35.00%, 9.01%, 44.60%, and 7.03%, respectively. These results indicate that TB-6 has a good repairing effect on the soil after solvent extraction, and the combined technology efficiently removed mechanical oil from the soil and reduced the damage caused by the solvent.

## Introduction

Mechanical oils are composed of 70% to 80% aliphatic hydrocarbons, 20% to 30% aromatic hydrocarbons, and a large number of volatile organic compounds and semi-volatile organic compounds (Ying et al. 2017). At a contaminated site, adsorption of mechanical oils to specific components of the soil hinders the natural degradation of the mechanical oils (Manilal and Alexander 1991).

Chemical treatments of sites contaminated by mechanical oils are mainly divided into two schemes: passivation treatment and removal treatment. The key to passivation treatment is to reduce the environmental pollution caused by pollution factors, for example, curing the migration of pollution factors with a curing agent. Previous studies on this scheme focused on the direction of heavy metals (Liu et al. 2018; Li et al. 2018). The removal treatment aims to change the influence of the pollution factor on the environment by using a redox reaction or leaching and extraction. The new substances generated by the redox reaction weaken or neutralize the toxicity of the soil, and leaching and extraction greatly reduce the concentration of the pollution factor in the environment; both methods can make the pollution factor less harmful or completely harmless to the environment (Tzaitang et al. 2009; Gómez et al. 2010; Jinlan et al. 2011). However, leaching and extraction methods damage soil sustainability, even when high-efficiency composite solvents are used and the process is optimized.

Biological treatments do not use chemical reagents, so secondary pollution does not occur. Although biological treatments have drawbacks, such as decreased processing efficiency, they enhance the self-purification ability of the soil and degradation by microorganisms or enrichment by plants that degrade or naturally consume mechanical oils (Khan et al. 2013).

In the soil, microbial degradation methods improve microbial population diversity, promote material circulation (He et al. 1997; Jim 2009), and reduce the impact of pollutants (Potthoff et al. 2006; Chandra et al. 2013) by the addition of exogenous microorganisms or strengthening of the original microorganisms. Therefore, a combination of chemical and biological treatments can not only increase scheme applicability but also greatly change the status quo of remediation of sites polluted by mechanical oils in China (Sutton et al. 2011).

In this study, we used a combination of solvent extraction and indigenous microbial degradation to effectively remove high concentration of pollutants and reduce the damage caused by the solvent in contaminated soil at an oil storage site. The original strain was isolated from paste sludge at the oil storage site, and the contaminated soil was used for domestication and acclimation of the strain. After the pollution factors were degraded using bio-enzymes and surfactants (Banat 1995; Pornsunthorntawee et al. 2008; Abbasian et al. 2015; Varjani and Upasani 2016), which are metabolized by the domesticated microorganisms, the soil quality and plant growth indicators before and after the treatment were compared to verify the repair effects of the combined technology on the contaminated soil. The soil quality and plant growth indexes were used to study the mechanism underlying soil remediation by solvent extraction–indigenous microbial degradation technology, and the high concentration of mechanical oil in the contaminated soil at an actual oil storage site was studied, which greatly improved the theoretical significance and practical application of this study.

## Materials and Methods

### Materials

#### Primitive soil

Dark brown soil was collected from 70–100 mm below the surface of the right side of a collection pool in the oil storage workshop of an assembly plant in Jiulongpo District, Chongqing City. The smell of mechanical oil was obvious during collection. The soil sample was sealed and stored in a self-sealing bag and air-dried in a fume hood in the laboratory for 72 h. Then, impurities such as stone particles were removed, ground into a powder, passed through a 0.425 mm sieve, and stored in a reagent bottle labelled S0. According to a third-party agency (Green Earth, Wuxi, Jiangsu Province, China), the mechanical oil content of the soil was 8.43%, of which lipids accounted for the largest proportion.

#### Preparation oil

According to the concentration and type of pollutants in the original soil and mass ratio of heavy-duty gear oil:waste hydraulic oil:diesel (5:3:2), *n*-hexane was used as the solvent to prepare simulated mixed soil for domestication of the microorganisms. All the raw materials for the media were purchased from Chongqing Laibosi Biotechnology Co., Ltd, Chongqing, China.

### Screening and domestication of indigenous microorganisms

#### Enrichment, primary screening, and re-screening

For enrichment, paste sludge from the oil storage site was inoculated into 100 ml of inorganic salt medium, placed in a constant-temperature shaking incubator, and incubated at 180 rpm and 28 °C for 5 days. For primary screening, 10 ml of the enriched culture solution was added to 250 ml Erlenmeyer flask of fresh inorganic salt medium (100 ml in each flask); heavy-duty gear oil, waste hydraulic oil, and diesel oil (0.5 ml each) were added to individual flasks and tested separately; the flasks were incubated at 180 rpm and 37 °C for 5 days; and this process was repeated 7 times. Then, the mixture was diluted, and 1 ml of the mixture was spread on nutrient agar plates and cultured at 37 °C for 24 h. The best-growing strain was selected and transferred to a sterile test tube containing nutrient agar. After primary screening, 0.5 ml of preparation oil was added to 10 ml of the culture solution and fresh inorganic salt medium and incubated at 180 rpm and 37 °C for 5 days; this step was repeated 7 times. The strain that showed good growth (measured for 48 h) and the highest degradation rate in 5 days was selected as the test strain and transferred to a sterile test tube containing nutrient agar.

#### Domestication of the test strain

The test strain was added to an Erlenmeyer flask containing the inorganic salt medium and mixed mechanical oil and cultured in a constant temperature water bath at 28 °C and 180 rpm for 24 h. The bacterial solution was used as the seed liquid. Next, 10 g of the simulated mixed soil was placed in a 250 ml Erlenmeyer flask; sterilized inorganic salt medium was added at a solid:liquid ratio of 1:10 and inoculated with 10 ml of the seed liquid; the flask was incubated at 37 °C and 140 rpm for 5 days; and the process was repeated 7 times to obtain a domesticated strain.

#### Identification of the domesticated strain

Submitted to a third party (Shanghai Majorbio Bio-pharm Technology Co.,Ltd.) for inspection.

### Solvent extraction of the contaminated soil

Using petroleum ether as the extraction solvent, the original contaminated soil, S0, was extracted twice by using 40 kHz ultrasonic power, 1:5 solid:liquid ratio, and 20 min extraction time, and the treated soil was labelled S1.

### Microbial inoculation of the contaminated soil

The domesticated strain was inoculated in nutrient agar and cultured for 18 h to prepare the seed liquid, 1 ml of which was inoculated into the degradation medium (0.5 ml of preparation oil was added to fresh inorganic salt medium) and cultured in the constant-temperature incubator for 24 h to obtain the culture solution. To prepare the solvent extraction–indigenous microbial degradation soil labelled S2, S1, culture solution, and deionized water (1:1:2) were incubated for 30 days in a custom-made ceramic pot (50 cm long, 15 cm wide, and 15 cm deep) with a lid.

### Soil quality test

The soil quality of S0, S1, and S2 was evaluated (Tian and Feng 2008; Harris 2010). The mechanical oil content in the soil was analyzed using gas chromatography-mass spectrometry (GC-MS); organic matter content in the soil, potassium dichromate method (Nelson et al. 1996; Ji 2005); water-holding capacity, centrifugation (Wang et al. 2007; Gao et al. 2010; Xing et al. 2016) and soil microstructure, scanning electron microscopy (SEM).

### Plant growth test

S0 and S1 were prepared (1:3 soil:deionized water) and placed (4 kg each) in planting pots; the lids were closed for 30 days. Then, alfalfa was planted in the pots containing S0, S1, and S2 (4 kg of soil in each planting pot). After 45 days, root and leaf growth of the plants was observed, and physiological and biochemical indexes of the plants were examined to evaluate the effects of the soil on the plants after solvent extraction–indigenous microbial degradation.

#### Root vitality

Determination of plant root activity by reducing triphenyltetrazolium chloride(TTC) method(Joslin and Henderson 1984; Yu 1992; Bat et al. 1994).

#### Plant protein content

Determination of plant protein content by using the Coomassie Brilliant Blue G250 Test method(Jones et al. 1989; Yu 1992; Seevaratnam and Patel BPHamadeh 2009) with a Bradford kit which was purchased from Chongqing Laibosi Biotechnology Co., Ltd, Chongqing, China.

#### Soluble sugar content

Determination of soluble sugar content by using anthrone colorimetry(Yu 1992).

#### Malondialdehyde (MDA) content

Determination of malondialdehyde (MDA) content by using thiobarbituric acid (TBA) to produce a color reaction with malondialdehyde in tissues under acidic conditions to produce reddish brown 3,5,5-trimethyloxazole 2,4-dione (trimethoate)(Hodges et al. 1999; Lykkesfeldt 2001).

#### Hydrogen peroxide content

Determination of hydrogen peroxide content by using method of ferrous oxidation of xylenol orange (FOX1 assay)(Bleau et al. 1998; Bou et al. 2008).

### Determination of mechanical oil content in the soil

#### Pretreatment

The Soxhlet extraction method was used to extract the mechanical oil from the soil samples. Acetone:hexane (1:1) was used for the extraction (Siddique et al. 2006; Saari et al. 2007), and petroleum ether was used as the extractant. The extraction solvent was evaporated to dryness under reduced pressure by using a rotary evaporator, and the material remaining after volatilization was dissolved in 10 ml of *n*-hexane to ensure uniform dilution for GC-MS.

#### Sample analysis

The pretreated samples were analyzed using GC-MS to determine the total concentration of the C10–C40 extract. HP-5 quartz capillary column (30 m × 0.25 mm × 0.25 μm) was used; the carrier gas was N_2_, and the temperature control range was 45–310 °C. The MS conditions were as follows: electron bombardment (EI) ion source; electron energy, 70 eV; transmission line temperature, 280 °C; ion source temperature, 230 °C; total ion scan mode; mass scan range (m/z), 60–640 amu (Yang et al. 2014).

### Statistical analysis section

All experimental data were set to 3 repetitions at least and analyzed by one-way analysis of variance (ANOVA) using a commercially available statistics software (OriginPRO for Windows, version 9.0; OriginLab Corp. Northampton, MA, USA). The results are expressed in the table as mean+SD. Values of p<0.05 were regarded as statistically significant.

## Results

### Screening and domestication of indigenous microbes

The heavy-duty gear oil, waste hydraulic oil, and diesel oil were separately screened to obtain 19 strains; then, the mixed mechanical oil was re-screened, and 6 strains were obtained (Fig. 1 and Table 1). The biomass of 6 strains reached OD_600_ > 0.3, i.e., CFU/ml > 3.0E+9, wherein the biomass of G-1, G-6, and H-6 exceeded 4.0E+9 CFU/ml (the precise biomass of H-6 was 3.98E+9 CFU/ml). The degradation rate of the 6 strains in 5 days was H-6 > G-3 > G-6 > H2 > H5 > G-1. Although the biomass values of G-1 and G-6 were higher than that of H-6, the degradation rate of G-1 was the lowest (1.63%) among the 6 strains, and the degradation rate of G-6 was much lower than that of H-6. Because H-6 showed the highest degradation rate (5.62%) and its biomass was higher than 4.0E+9 CFU/ml, it was selected for domestication and labelled as Test Bacteria-6 (TB-6).

**Fig. 1.** Biomass of strains during rescreening Note: Biomass are given as the mean ± SD for 3 Erlenmeyer flask of culture solution in each group. Pictures are produced by OriginPRO after analyzing data

**Table 1.**
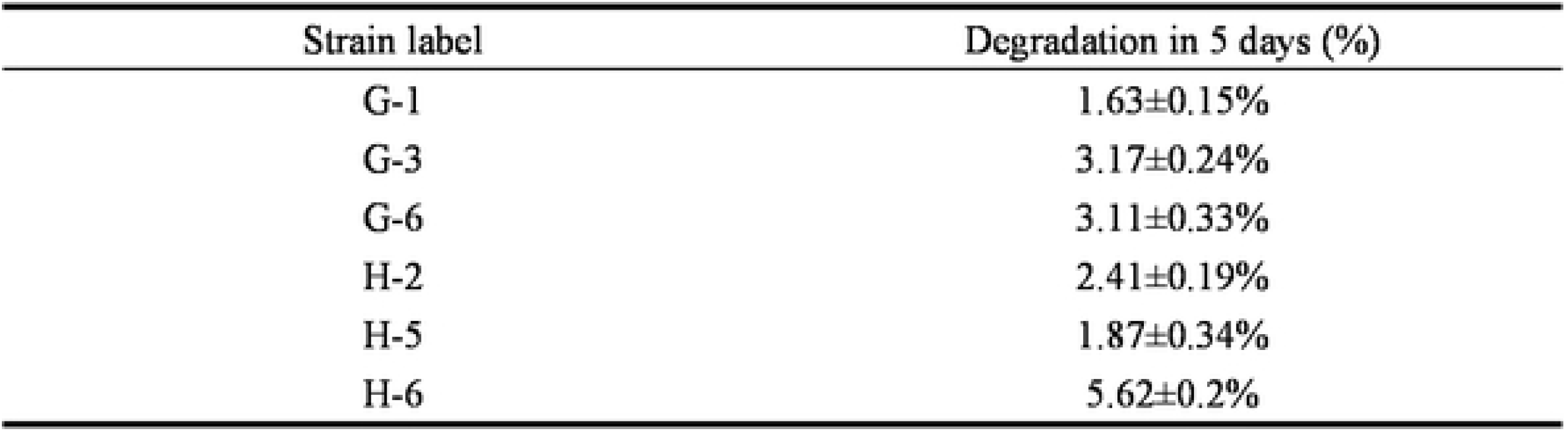
Degradation rate of strains during 5 days of rescreening Degradation data are given as the mean ± SD for 3 Erlenmeyer flask of in each group.

| Strain label | Degradation in 5 days (%) |
| --- | --- |
| G-1 | 1.63±0.15% |
| G-3 | 3.17±0.24% |
| G-6 | 3.11±0.33% |
| H-2 | 2.41±0.19% |
| H-5 | 1.87±0.34% |
| H-6 | 5.62±0.2% |

Using the original strain H-6 as the control, the domesticated strain was inoculated in nutrient agar, cultured for 48 h to prepare the seed liquid and 10 ml of the seed liquid was mixed with the inorganic salt culture solution containing mixed mechanical oil and cultured for 5, 10 and 15 days. The effects of domesticated strain TB-6 on the degradation rate of mixed mechanical oil and segmentation of the petroleum hydrocarbon carbon chains were evaluated (Table 2 and Fig. 2).

**Table 2.**
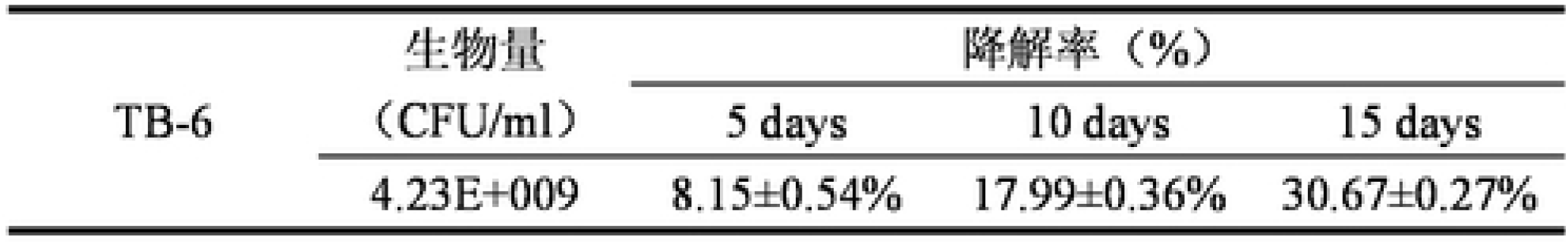
The biomass and degradation rate of the domesticated strain Degradation data are given as the mean ± SD for 3 Erlenmeyer flask of culture solution in each group.

**Fig. 2.** Variations in mixed mechanical oil content with segmentation of the petroleum hydrocarbon chains Note: Values from the detection report of Analysis and Testing Center of Chongqing University, Chongqing, China

The biomass of strain TB-6 increased by 6.3% after acclimation. The degradation rate of mixed mechanical oil was 8.15%, which was 46.26% higher than that before acclimation, and the degradation of mixed mechanical oil increased as the degradation time increased (17.99% in 10 days and 30.67% in 15 days; Table 2). After 15 days, the total content of the mixed mechanical oil decreased from 531.7 mg/g to 368.6 mg/g when compared with the control group, and the degradation rate was 30.67% (Fig. 2). The carbon chain length of C15–C28 decreased by 24.9%, i.e., 132.4 mg/g and that of C29–C40 by 7.52%, i.e., 40.0 mg/g and that of C10–C14 increased by 1.75%, i.e., 9.3 mg/g. In summary, the domestication of indigenous microorganisms was effective. The degradation rate of mixed mechanical oil by TB-6 increased as the degradation time increased, and TB-6 mainly degraded the medium- and long-chain petroleum hydrocarbons.

### Characteristics of the TB-6 strain

The morphological characteristics of the TB-6 strain are listed in Table 3. The colonies are small, round, and milky white with a smooth surface, moist luster, neat edge, and no rough branches. The whole body is translucent. After Gram staining, obvious pink bacteria can be observed under the microscope. The single granule of the bacteria is dispersed, and short rods can be mostly observed.

**Table 3.** Morphological characteristics of the TB-6 strain

| Strain | Colony Characteristics |  |  |  |  |  | Gram staining | Microscopic characteristics |
| --- | --- | --- | --- | --- | --- | --- | --- | --- |
|  | Size | Form | Color | Status | Border | Transparency |  |  |
| TB-6 | Small | Circular | Milky white | Smooth moist | Neat | Translucent | G (-) | Single grain short rod |

The physiological and biochemical results of strain TB-6 are listed in Table 4. Enterobacter TB-6 is a facultative, fermentative anaerobic gram-negative bacterium, and the V-P and indole tests yielded negative results. TB-6 cannot hydrolyze starch, but it can produce H_2_S. It can use citric acid and has a hydrogen peroxide catalyst. Methyl red, oxidase, and nitrate reduction tests yielded positive results. The optimum growth pH of the strain is 6.5–7.0, and the optimum growth temperature is 36–39 °C. The accession number for TB-6 in the GenBank database is MK509013.

**Table 4.** Physiological and biochemical characteristics of the TB-6 strain Note: +, positive; −, negative.

| Strain | Parameter | Performance |
| --- | --- | --- |
| TB-6 | Oxygen tolerance | Facultative anaerobic |
|  | Acid and alkali tolerance | 6.5–8.5 |
|  | Growth temperature | 27–41 °C |
|  | Types of oxidative fermentation of sugars | Fermentation type |
|  | NaCl tolerance | 9% |
|  | Mobility | + |
|  | V-P | - |
|  | MR | + |
|  | Hydrolysis ability of starch | - |
|  | Citrate utilization ability | + |
|  | Indole | - |
|  | Hydrogen sulfide production ability | + |
|  | Nitrate reduction ability | + |
|  | H<sub>2</sub>O<sub>2</sub> catalyst | + |
|  | Oxidase | - |

### Effects of solvent extraction–indigenous microbial degradation on mechanical oil content in the soil

The mechanical oil contents in soils S0 and S1 before and after solvent extraction are shown in Fig. 3. The mechanical oil content in S0 was 84.3 mg/g, mechanical oil content in S1 after solvent extraction was 2.5 mg/g, and mechanical oil removal rate was 97.03%.

**Fig. 3.** Analysis of mechanical oil content in the soil before and after solvent extraction by using GC-MS Note: Figure from the detection report of Analysis and Testing Center of Chongqing University, Chongqing, China

After degradation, the maximum intensity of mechanical oil content in S2 was 0.68%, 10.40%, and 19.45% lower than that in S1 in 5, 15, and 30 days, respectively (Table 5). The difference in mechanical oil content led to a significant (P<0.05) decrease between 15 and 30 days than between 5 and 15 days, indicating that the mechanical oil content in the soil was further reduced with the increase in degradation time (Fig. 4).

**Table 5.** Maximum intensity of GC-MS peaks in mechanical oil content in soil S2 Values from the detection report of Analysis and Testing Center of Chongqing University, Chongqing, China

| Soil label | S1 | S2 |  |  |
| --- | --- | --- | --- | --- |
| Maximum intensity | 46,370,403 | 5 days<br>46,051,915 | 15 days<br>41,549,396 | 30 days<br>37,352,158 |

**Fig. 4.** GC-MS detected changes in the mechanical oil content in S2 Note: Figure from the detection report of Analysis and Testing Center of Chongqing University, Chongqing, China

### Effects of solvent extraction–indigenous microbial degradation on soil microstructure

The organic matter content in the soil was in the order of S0 ≫ S1 < S2 (Fig. 5), and the soil water-holding capacity was in the order of S0 < S1 < S2 (Fig. 7). S0 had an agglomerate structure, the knot was obvious, water-holding capacity of S0 was only 31.37% (because drying treatment increases the volatility of the original pollution factor, the actual water-holding capacity of S0 may be lower than the measured value) and large fluctuations were observed during data processing. After extraction with petroleum ether, most of the mechanical oil content in S1 was removed, so the water-holding capacity of S1 was 68.20% higher than that of S0. However, the organic matter content in S1 was reduced by 82.98% when compared with S0. The soil microstructure is shown in Figure 6; the S1 agglomerate was greatly reduced but more compact. After inoculation of S2 with TB-6 and 30 days of natural growth, the organic matter content in S2 increased by 35.00% when compared with S1, and the surface of S2 was rough with loose particles (Fig. 6). The water-holding capacity of S2 was 9.01% higher than that of S1 and 83.35% higher than that of S0 (as shown in Fig. 8).

**Fig. 5.** Comparison of organic matter contents in the different soils Note: Values from the detection report of Green Earth, Wuxi, Jiangsu Province, China

**Fig. 6.** Comparison of microstructures of the different soils

**Fig. 7.** Comparison of water-holding capacities of the different soils Note: Values are expressed as mean ± SD for 3 groups of soil in each group. Pictures are produced by OriginPRO after analyzing data

**Fig. 8.** Comparison of the appearance of alfalfa grown in the different soils

### Effects of solvent extraction–indigenous microbial degradation on plant growth

#### Root and leaf growth

The total length, chlorophyll content, nitrogen content, and root growth of alfalfa were in the order of S0 < S1 < S2 (Fig. 8 and Table 6). Forty-five days after planting, the growth of roots and leaves in S0 was lower than the growth of roots and leaves in S1 and S2. The alfalfa in S2 showed the best growth.

**Table 6.**
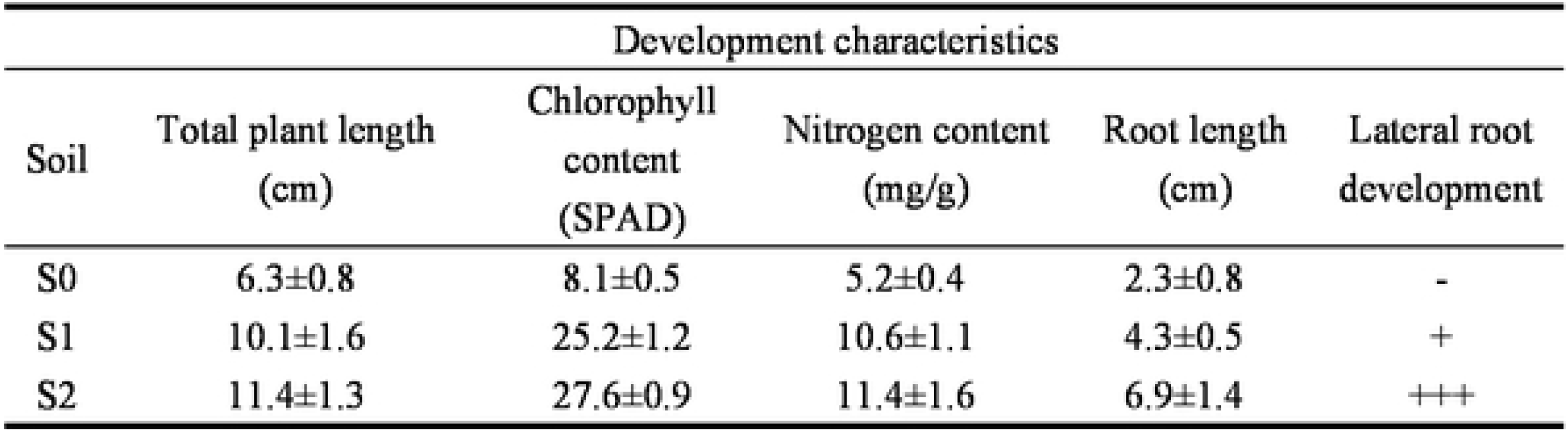
Whole plant development Note: −, undeveloped; +, developed with good density; ++, excellent density. Values are given as the mean ± SD for 5 plants in each group.

| Development characteristics |  |  |  |  |  |
| --- | --- | --- | --- | --- | --- |
| Soil | Total plant length<br>(cm) | Chlorophyll<br>content<br>(SPAD) | Nitrogen content<br>(mg/g) | Root length<br>(cm) | Lateral root<br>development |
| S0 | 6.3±0.8 | 8.1±0.5 | 5.2±0.4 | 2.3±0.8 | - |
| S1 | 10.1±1.6 | 25.2±1.2 | 10.6±1.1 | 4.3±0.5 | + |
| S2 | 11.4±1.3 | 27.6±0.9 | 11.4±1.6 | 6.9±1.4 | +++ |

#### Physiological and biochemical indexes of the plants

##### Root vitality

In the plants, root vigor was in the order of S2 > S1 > S0 (Fig. 9a); plant root activity significantly(P<0.05) increased by 44.60% in S2 when compared with S1, and it about 2.5 times significantly(P<0.05) higher than that in S0.

**Fig. 9.** Physiological parameters of alfalfa in the different soils: (a) root vigor (b) plant protein content (c) soluble sugar content (d) MDA content (e) HP content Note: Values are expressed as mean ± SD for 3 groups of approximate samples in each group. Pictures are produced by OriginPRO after analyzing data.

##### Plant protein content

In the plants, protein content detected using the Bradford method was in the order of S2 > S1 > S0 (Fig. 9b); the total protein content of the plants in S2 increased by 0.083±0.007μg/μl when compared with S1, which was higher than that of the plants in S0 (0.1871±0.021μg/μl).

##### Soluble sugar content

In the plants, soluble sugar content was in the order of S0 > S2 > S1 (Fig. 9c), and the soluble sugar content of the plants in S0 was 39.16% and 30.00% significantly(P<0.05) higher than that in S1 and S2, respectively. The soluble sugar content of the plants in S2 was 7.03% significantly(P<0.05) higher than that of the plants in S1.

##### Malondialdehyde (MDA) content

In the plants, MDA content was in the order of S2 > S1 > S0 (Fig. 9d), and MDA content in the plants in S2 was 68.68% significantly(P<0.05) higher than that in S1 and about 2.25 times significantly(P<0.05) higher than that in S0. The soil treatment increased the MDA content of the plants.

##### Hydrogen peroxide content

In the plants, hydrogen peroxide concentration was in the order of S0 > S2 > S1 (Fig. 9e).

## Discussion

### Effects of solvent extraction–indigenous microbial degradation on the soil

During the extraction process, Because of the “like-dissolves-like” rule (Montes 2003) between the extraction solvent petroleum ether and mechanical oil in the soil, the mechanical oil in the soil enters the solvent liquid phase and is mostly separated and removed after the extraction solvent is added to the contaminated soil S0. However, because of the same rule, the extraction solvent has a huge impact on the organic matter content in the soil, resulting in damage to soil quality. With respect to microbial degradation, the mechanical oil available for TB-6 was reduced (Pignatello and Xing 1996; Luo et al. 2004), and the bioavailability was decreased (Tabak and Govind 1997). On the basis of solvent extraction, the bioavailable degradation rate of strain TB-6 in the soil of the oil storage site reached 27.4% in 50 days. The solvent extraction–original microbial degradation technology was used for the total mechanical oil content of the oil storage site. The removal/degradation rate was 97.85%, and the residual mechanical oil content in S2 soil was 1.815 mg/g.

After extraction by the organic solvents, most of the soil particles were unsupported and soil bulk density increased, which caused soil consolidation in S1. This phenomenon is consistent with that reported by previous studies (Zhuang et al. 1999; Xu and Wang 2005; Nawaz and Trolard 2013). Although the organic soil content in S0 was very high (Fig. 6), the molecular action of petroleum hydrocarbons caused the aggregate structure to become tightly combined to form a knot; moreover, the internal structure was unstable and easily deformed by the petroleum hydrocarbon particles. After the inoculation of S1 by TB-6, the organic matter in the soil increased (Fig. 5), and the material support between the soil particles was supplemented by the action of microorganisms. Simultaneously, the soil structure became loose because of the reduction in the number of petroleum hydrocarbon particles, and the surface structure of the pellets became rougher.

The oil-containing aggregate structure of S0 showed reduced hydrophilicity, which resulted in decreased water-holding capacity of S0 and data fluctuations. After the extraction, the hydrophobic layer disappeared, and the water-holding capacity of S1 was restored. Because microbial inoculation restores the organic matter content through the metabolic behavior of the microorganism, the interstitial space between the soil particles in S2 was restored and the S2 surface was more crisp and rough (as shown in Fig. 6), increasing the water-holding capacity of the soil aggregate structure (Mapa 1995; Rawls et al. 2003; Haverkamp et al. 2005).

### Effects of solvent extraction–indigenous microbial degradation on the plants

In S0, root growth and development were inhibited because of the presence of petroleum hydrocarbons (Clarke and Ward 1994); the reduction of 1% 2,3,5-triphenyltetrazolium chloride was low (Fig. 8), which is consistent with the results of (Li et al. 1997; Ravanbakhsh et al. 2009). After solvent extraction, the roots in S1 developed rapidly when compared with those in S0 because the hydrophobic layer formed by the mechanical oil on the soil aggregate was eliminated, which restored the normal material exchange between the soil and plant. Therefore, alfalfa growth was better in S1 than in S0. However, the extraction resulted in the loss of soil organic matter and increased soil bulk density and compaction (Zhuang et al. 1999; Xu and Wang 2005). Such knots inhibit root growth (Nadian et al. 2010) and make it difficult to maintain nutrient circulation after root growth because the organic matter content in the solvent-extracted soil is low; thus, root activity was higher in S2 than in S1. After the microbial treatment, the plant growth in S2 improved further because of the effects of microbial remediation. This also confirms the findings of Li et al. (1997), who reported that the soil-water relationship is one of the most important factors for assessing plant growth in bioremediated soils. Our findings show that the inoculated microorganism promoted root activity in the plants, which may be attributable to the following 2 aspects:

1. The microorganisms degrade the mechanical oil remaining in the soil after solvent extraction, so inhibition of the growth and development of plant roots by the mechanical oil is weakened. The use of microorganisms to degrade petroleum hydrocarbons reduces the uptake of petroleum hydrocarbons from the soil during the growth and development of plant roots.
2. The physiological activities of microorganisms cause plant roots to be vigorous. Microorganisms inevitably produce metabolites, and these products stimulate the development of plant roots. (Napora et al. 2014) and (Zhou et al. 2015) have reported the promotion of root growth by microorganisms.

Small molecules of petroleum hydrocarbons may affect plant genes after the plants absorb petroleum hydrocarbons (Pasquevich et al. 2013). Furthermore, protein synthesis was slowed down or changed, resulting in less accumulation of proteins in the S0 environment than in the S1 and S2 environments. Solvent extraction reduced the amount of petroleum hydrocarbons in the soil, restored the growth of plant roots in S1 and S2, and enhanced the ability of the plants to obtain nutrients from the soil. Because of the intervention of TB-6, to some extent, the organic matter lost in S1 was supplemented and nitrogen circulation in the plants was improved. Therefore, the amount of protein synthesis in the plants in S2 was slightly improved when compared with S1 (Clement and Gessel 1985).

In plants, soluble sugar is the main product and energy storage form of photosynthesis (Xia et al. 1982), and it helps during cold resistance (Wang 1996; Wang et al. 1998) and regulates physiological activities (Zhao et al. 2006). The plants in S0 had a much higher soluble sugar content after solvent extraction because water circulation through the root system is hindered, directly increasing the liquid concentration outside the plant cell. Therefore, soluble sugar content will increase in the plant to maintain the osmotic pressure and ensure normal operation of the physiological system (Yamane and Ichiro 1969; Zhang et al. 2016). Improvement of soil organic matter in S2 increased the quality of substances entering the plants and concentration of extracellular liquid, resulting in an increase in the soluble sugar content.

MDA reflects the degree of peroxidative damage to the plant tissue or membrane lipids of plant organelles, which is inconsistent with the plant growth shown in Figure 8. The reasons for such anomalies may be as follows:

1. Because organic solvent extraction removes most of the organic matter from the soil, the soil quality becomes poor. Simultaneously, soil compaction is observed, causing environmental stress on the plants and increasing the MDA content (Liu and Wu 2010).
2. The physiological activities of the inoculated microorganisms induce an allelopathic effect on the plants (Molisch 1938; Rice 1983) and cause them to have a stress response to the environment and peroxidation of cell membranes and further increase in MDA content. Although this allelopathic phenomenon caused an increase in MDA content, it also promoted the growth of alfalfa. (Hasegawa et al. 1992) and (Li 1989) have reported the promoting effects of the allelopathic phenomenon in plants.
3. The level of plant development was lower in S0 than in S1. In addition, the plants in S2 grew better than those in S1 because of the effects of the microorganisms. Therefore, differences in the MDA contents may be attributable to the differences in plant growth levels in the 3 types of soils.

In S0, the petroleum hydrocarbons caused obvious stress to the plants, which resulted in increased accumulation of hydrogen peroxide (Gechev and Jacques 2005; Ireneusz et al. 2007). After solvent extraction, the hydrogen peroxide content in the plants in S1 was reduced by 10.07% when compared with S0. The environmental stress induced by TB-6 increased the hydrogen peroxide content slightly in the plants; however, in general, hydrogen peroxide content in alfalfa in S2 was normal. Therefore, only a slight fluctuation was observed between the hydrogen peroxide contents in S1 and S2.

## Conclusions

The purpose of this study was to use an effective chemical–biological treatment scheme to efficiently remove high concentrations of mechanical oil from contaminated soil and reduce the damage caused by chemicals to soil quality. This study mainly consisted of three parts, i.e., screening and domestication of in situ microorganisms, analysis of the effects of solvent extraction–indigenous microbial degradation on soil quality, and evaluation of the effects of solvent extraction–indigenous microbial degradation on plant growth.

The solvent extraction removed 97.03% of the mechanical oil in the contaminated soil, reduced soil agglomeration, increased water-holding capacity by 68.20%, and restored the material exchange capacity between plants and soil. It also enhanced root activity and soluble sugar, protein, and hydrogen peroxide contents in the alfalfa plants. However, 82.98% of the organic matter in the soil was depleted, making the soil more compact and affecting soil quality. Although solvent extraction helps to remediate soils contaminated by mechanical oils, it also causes great loss to soil organic matter; therefore, inoculation of microorganism could be used to ameliorate the adverse effects of solvent extraction on contaminated soils.

The solvent-extracted soil was inoculated with TB-6 for 30 days; the degradation rate was 19.45%. For 50 days the mechanical oil content in the soil decreased to 1.815 mg/g and The bioavailability was stable at about 27.4%. This means the solvent extraction–indigenous microbial degradation technology removed/degraded mechanical oil from the contaminated soil at a rate of 97.85% at last. The strain made the soil particles after solvent extraction loose and rough, soil organic matter content increased by 35.00%, water-holding capacity increased by 9.01%, root activity of alfalfa increased by 44.60%, plant protein content increased by 0.0198 mg/ml, and soluble sugar content increased by 7.03%. This indicates that the strain TB-6 has a good repairing effect on the contaminated soil after solvent extraction.

Inoculation of the soil with TB-6 induced the allelopathic effect on the alfalfa plants, and the MDA content of the plants increased further. To ensure that soil function is not affected, it is necessary to study the allelochemicals produced by TB-6.

